# A possible connection between vertical transhumance and temporary plague reservoirs during the Second Plague epidemic

**DOI:** 10.1101/2025.10.10.681562

**Authors:** Nimrod Marom, Jörg Baten, Philip Slavin

## Abstract

The Second Plague Pandemic (SPP) ravaged large parts of Eurasia and North Africa between the 14th and 19th centuries. Today, plague is still active in parts of Asia, Africa, and America. One of the hotly debated topics in plague studies is the geographic origins of the recurrent plague waves in Europe after the Black Death (1338-1353): Were they being repeatedly introduced from ‘outside’, or did they originate in ‘domestic’ reservoirs? Here, we build upon recent work arguing for the existence of domestic reservoirs, by exploring a possibility that vertical transhumance by pastoralists could have been a conduit between temporary plague reservoirs and human population centers in Europe. We argue that aspects of pastoral movement and practice, including the utilization of marginal areas and of caves, account for some aspects of plague outbreaks both in the present and in the past. We support this hypothesis using historical sources from the Second Plague Pandemic in Central Europe, and suggest an association between recent and historical land use for pasture and plague outbreaks data.

## 1 Introduction

The Second Plague Pandemic (SPP) originated in Central Asia in the late 1330s with what became known as the Black Death (1338-1353) and has recurred in dozens of subsequent waves from the mid-fourteenth to the early nineteenth century. The SPP had a profound impact on different demographic, socio-economic, cultural and political processes on a quasi-global level (1, 2). Although effectively extirpated from Europe, plague-related morbidity and mortality still occur in Africa, Asia, and America, with World Health Organization reports listing a few hundred cases each year (3, 4). Epidemiological studies of the plague are still ongoing (4–8), with the overarching goal of preventing new outbreaks and monitoring the emergence of antibiotics-resistant strains (9).

The plague is a zoonotic disease, caused by the bacterium Yersinia pestis, which has evolved multiple lineages and strains (10–12). The predominant reservoir species for plague are wild rodents (13, 14), while ectoparasites (mainly fleas) constitute the main vector for bacterial transmission from rodents to other mammals, resulting in bubonic plague. Septicemic and gastroenteritic forms of plague can be contacted by ingesting the meat of an infected animal, while pneumonic plague is being transmitted via inhaling infected aerosols (15). This basic scheme of rodent → flea → human Y. pestis transmission, however, is in practice determined by cultural processes and practices that form the epidemiological highway leading from rodent reservoirs to human populations.

The pandemic aspect of the SPP has been famously associated with the ubiquity of its commensal rodent species, the black rat (Rattus rattus), and this rodent’s ability to spread along maritime trade routes (16, 17). Thus, in many instances, plague would be imported from one region into port cities of another region, and then spread inland along river valleys (18, 19). However, maritime transportation of flea-infested, plague-carrying rats is but one of many biocultural pathways transmitting Y. pestis from reservoirs into contact with human populations. Other biocultural pathways include warfare, with soldiers transmitting the bacteria either through direct contact or by infected ectoparasites (20); and trade, involving transportation of infested commodities, especially clothing, skins and furs that moved along with vectors such as fleas and human body lice (21, 22).

In the present paper, we explore yet another, hitherto understudied biocultural pathway, that of pastoral transhumance, as an enabler and catalyst of plague outbreaks in inland locations. The potential role of nomadism for contacting and spreading plague in past centuries has been briefly touched upon in the context of the Ottoman Empire, questioning the previous view that low population densities and mobile lifestyle contributed to lower mortality rates in nomadic communities (21), and Central Asia at the onset of the Black Death (23, 24). However, the idea that pastoral transhumance has been a major biocultural pathway for spreading the Second Plague from local reservoirs to population centers in Europe has not been raised, to the best of our knowledge. We propose a potential link between transhumant pastoral activity and plague outbreaks, and examine this hypothesis with a historical analysis of contemporary and Second Plague outbreak datasets. Given that maritime and overland trade are no longer considered relevant for present-day plague occurrences, while pastoral transhumance persists, understanding this mechanism could be relevant for controlling current morbidity.

Pastoralism is intimately related to animal husbandry practices involving a substantial degree of seasonal mobility. Transhumance, the more precise term for tracking and securing food and water for livestock, helps pastoralists avoid the competition over land with farmers, and allows to maintain larger herds in challenging environments by utilising seasonal plenty (25, 26). Challenging environments with abundant pastoral resources are a context-specific definition. While in North Africa and the Middle East seasonal water availability makes desert edges a viable option for pastoral transhumance, in Europe vertical transhumance to upland summer pastures has been and is still being practiced (27, 28). Pastoral transhumance occurs on many scales, from herds numbering thousands of animals covering yearly trajectories hundreds of miles long, to small flocks grazing in regions a few kilometers away from the permanent settlement (29). The viability of different transhumant systems depends on environmental carrying capacity, seasonality, and demand for herd products leading to regional production specialisms (30, 31). However practiced and on whatever scale, transhumant pastoralism forms a potential biocultural pathway for facilitating plague spillover (zoonosis) to and outbreaks in humans. Empirical studies for the association of pastoral nomads with plague have taken place in 20th-century North Africa and Central Asia (32, 33), and emphasized the role of interaction of such human communities with wild rodent reservoirs (jirds and marmots, respectively) in propagating the disease.

Pastoralists are specifically prone to exposure to plague reservoirs due to their association with husbanded animals, some of which in European contexts - sheep, goats and dogs - are potential carriers of the disease, acting as intermediary hosts between wild rodents and humans (34–36). Specific mechanisms of transition from livestock and herd dogs to humans include flea bites, but also specific and ubiquitous activities such as butchery (7) or the consumption of infested meat (37). In addition, the geographical liminality of transhumant pastoralists, who visit remote regions on a yearly basis, makes them more likely to interact directly (through hunting and processing) or through vectors (through cohabitation and flea crossover) with wild animals that maintain plague reservoirs. Furthermore, the vertical transhumance phenomenon, characteristic of highland environments, could be yet another factor bringing herders and their flocks closer to natural reservoirs of the plague: as evidence from Central Asia, Caucasus and Western China indicates, the majority of reservoir maintaining rodents tend to thrive at higher altitudes (38–40), while pastoralists’ annual descend from the same altitudes at the end of transhumance season may be instrumental in facilitating the downward spread of plague into human habitat. Finally, the tendency of pastoralists to repeatedly occupy seasonal shelters during their transhumant rounds may be another potential factor creating temporary reservoirs, where the bacteria survive in sediments and local hosts to reinfect a flock once it returns to the same place. This is especially true for caves, constituting ecological nexuses where wild and domestic animals interact with each other on the one hand, and with humans on the other, in different ways (41–43).

The comfortable climate inside caves during seasonal extremes make these places home to rodents, ursids, hyraxes, and other potential reservoir species for Y. pestis (14), as well as bats and other mammals maintaining reservoirs of other pathogens (44). The biogenic cave sediment can both sequester ectoparasites acting as vectors (45), and serve as temporary reservoirs of bacteria, which, in turn, can survive in a virulent state in sediments for at least a year (46). Soil infection could derive from aerosols created by carriers or by decomposing cadavers (47). In the Middle East and much of Europe, use of cave entrances as seasonal shelters and pens in remote regions is ubiquitous. Caves are favoured as facilities of natural shelter, and for their agreeable temperature that averages the yearly extremes of heat in desert environments, and of cold in Alpine ones.

The use of caves to shelter herd animals is a trivial feature of karstic regions that it is rarely noted in the literature; Hammer’s (48) description of the situation in southeast Turkiye is typical of other regions (e.g., 49, 50, 51). Penning animals in caves where wild rodents and their fleas inhabit could cause infection among the livestock, which can also carry infected fleas down to populated regions towards the end of summer, when the flocks return to their winter pastures (Figure 1). The probability-focusing role of caves in space, constituting a small biological nexus where multiple organisms meet and where local ecologies created excellent conditions for trans-seasonal survival of the bacteria, repeatedly visited by shepherds with their livestock and dogs, may have been a crucial aspect of plague transmission first to these domesticated animals and then of its eventual spillover to humans and subsequent spread into lowland human populations.

**Figure 1:**
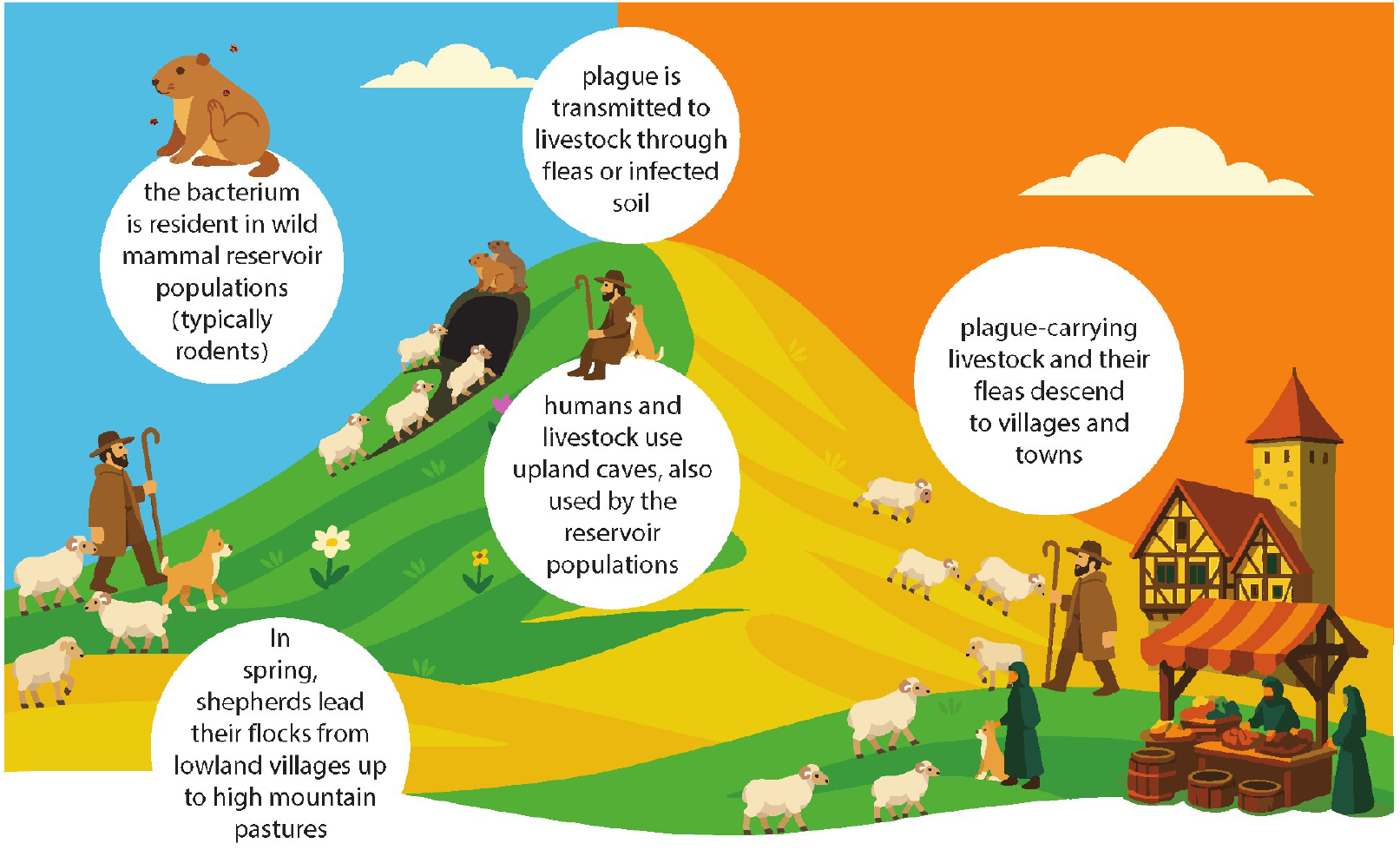
Sketch of the transhumant cycle, illustrating the correspondence between changes in elevation, the pastoral transhumance cycle, and the autumn outbreaks. Caves are suggested to have played a major role as a nexus where wild reservoirs, vectors, and domesticates meet. Drawing by Amal Hibner.

Transhumant pastoralism may thus establish a vertical ‘epidemic highway.’ This pathway would link wild reservoirs in high-elevation areas, used for summer grazing, with more populated lowland areas serving as winter pastures. We will investigate the correlation between herding intensity and plague occurrence using historical and contemporary data on plague outbreaks and pastoralism. Our historical analysis will concentrate on Central Europe, a region rich in historical records, enabling a detailed examination of the connection between transhumant herding practices and plague outbreaks.

## 2 Results

Although nomadism, in the colloquial sense, was from time to time associated in European contexts with the spread of contagion (as it happened in 1383 in Transylvania) (52), the association between the two variables (and plague in particular) was much more commonplace in the Islamic regions of North Africa and the Ottoman Empire (21). However, large-scale pastoral transhumance was no less common in European upland environments than those of the Middle East during the Second Plague Pandemic, from the mid-14th to the early 19th centuries CE.

A number of special characteristics were shared by the regions in which plague outbreaks were occurring in Southern Germany and some of which were situated closer to South German plague reservoirs. These regions were geographically situated in the Swabian and Franconian Jura mountains, the Southern Blackwood Forest, the Bavarian and the Bohemian Forest, and the Ore Mountains between Saxony and Bohemia (today’s Czech Republic), and the river landscapes next to these mountainous regions (53). This characteristic of mountainous terrain near populated river landscapes was important for the widespread use of vertical transhumance. During the spring and summer, sheep were brought to higher altitudes (mostly April to August) so that the fertile and densely populated landscapes along the rivers Neckar, Rhine, Danube, and others could be intensively used for grain and vegetable production. The system was originally introduced by the monastic reform order of the Cistercians, who encouraged peasants to substantially increase sheep herding. The Cistercians introduced a number of economic innovations in the late medieval period, and wool production was one of their most important projects in Southern Germany (54). In doing so, they set up an export-oriented textile production in Southwestern Germany, which provided woolen textiles in the 16th and 17th centuries to export markets as far as Poland, Northern Italy and France (55). For this, local wool production needed to be intensified. Although the large-scale vertical transhumance of sheep only started in the 18th century in Southwestern Germany, the strong intensification of sheep holdings by the peasants and lay brothers of the Cistercians already increased the contact between humans and animals since the late medieval period.

Southwestern Germany was also the most geographically mobile society of Central Europe. It was in that context that the Duke of Württemberg, was forced in the famous “Treaty of Tübingen” in 1514 to allow geographic mobility which was largely unconstrained in his territories, a unique feature within Central Europe (56). It was also in this wider geographic context of southern Germany that some late-medieval plague reservoirs were situated, as noted above.

Finally, these specific regions were the ones in Central Europe in which sparsely populated mountain regions and very densely populated fertile river regions were situated close to each other, making vertical transhumance most economically productive due to the short distance (57). The large demand for wool by the important textile production, not only in Southwestern Germany but also in Bohemia, Saxony, East Franconia and Northern Switzerland further encouraged herders and peasants to increase their sheep stocks. All these factors together, including the intensification of wool production in the late-medieval period, the commercialisation of textile industry in the early modern era, the proximity between densely populated fertile valley and sparsely populated mountain regions, and the geographic mobility of the population were all preconditions for vertical transhumance and a close contact between humans and animals. Two important features of the Second Plague Pandemic can be explained by applying the transhumant pastoralism hypothesis to the transmission of plagues from inland reservoirs to population centers in Europe. The first is the inland outbreaks of the disease, which are hypothesised to have been caused by the engagement of humans, livestock and commensal animals near short- or medium-term reservoirs. Although the question of plague reservoirs in Europe has been long debated by historians, it was only recently that a strong case for their existence has been made, based on empirical evidence, textual and aDNA as it was in the case during the pestis secunda wave commencing in South-Central Germany, possibly in southern Hesse, in 1356 (e.g., 58). Such inland commencement and initial spread of plague waves in Europe are typically explained by contingency, like the unspecified, random contact with infested goods or persons (18), or with wild rodents. The vertical transhumance as a biocultural transmission pathway, however, goes beyond contingency to provide a mechanism for connecting potential high-elevation reservoirs in Europe to lowland population centers (59).

Another feature of some of the Second Plague Pandemic outbreaks in Europe is their proclivity, in many instances, to occur in a period ranging from mid-summer to early autumn, as it is indeed revealed in parish burial records in early modern German-speaking lands (19). Likewise, summer outbreaks were often documented in other contexts of the Second Plague Pandemic (60), although plague might invade any given region at virtually any point during the year. This is perhaps somewhat surprising in the context of epidemic outbreaks, which should be more easily transmitted in enclosed spaces during winter; summer outbreaks, however, fit well with the annual seasonality of large-scale transhumance itineraries practiced in Central Europe. The Swabian-Franconian transhumant system consisted of the practice, whereby shepherds would move between extensive upland regions in the Swabian and Franconian Jura mountains and lowland wintering grounds sometimes tens or a couple of hundred kilometers away. The key calendar dates for the spring upland transhumance was 23rd of April (St. George’s Day), when the shepherds usually arrived in their summer grazing areas, and the 24th of August (The Feast of St. Bartholemew) when they again left on their way to their permanent homes at lower altitudes (61), often simultaneously with reported plague outbreaks (Figure 2).

**Figure 2:**
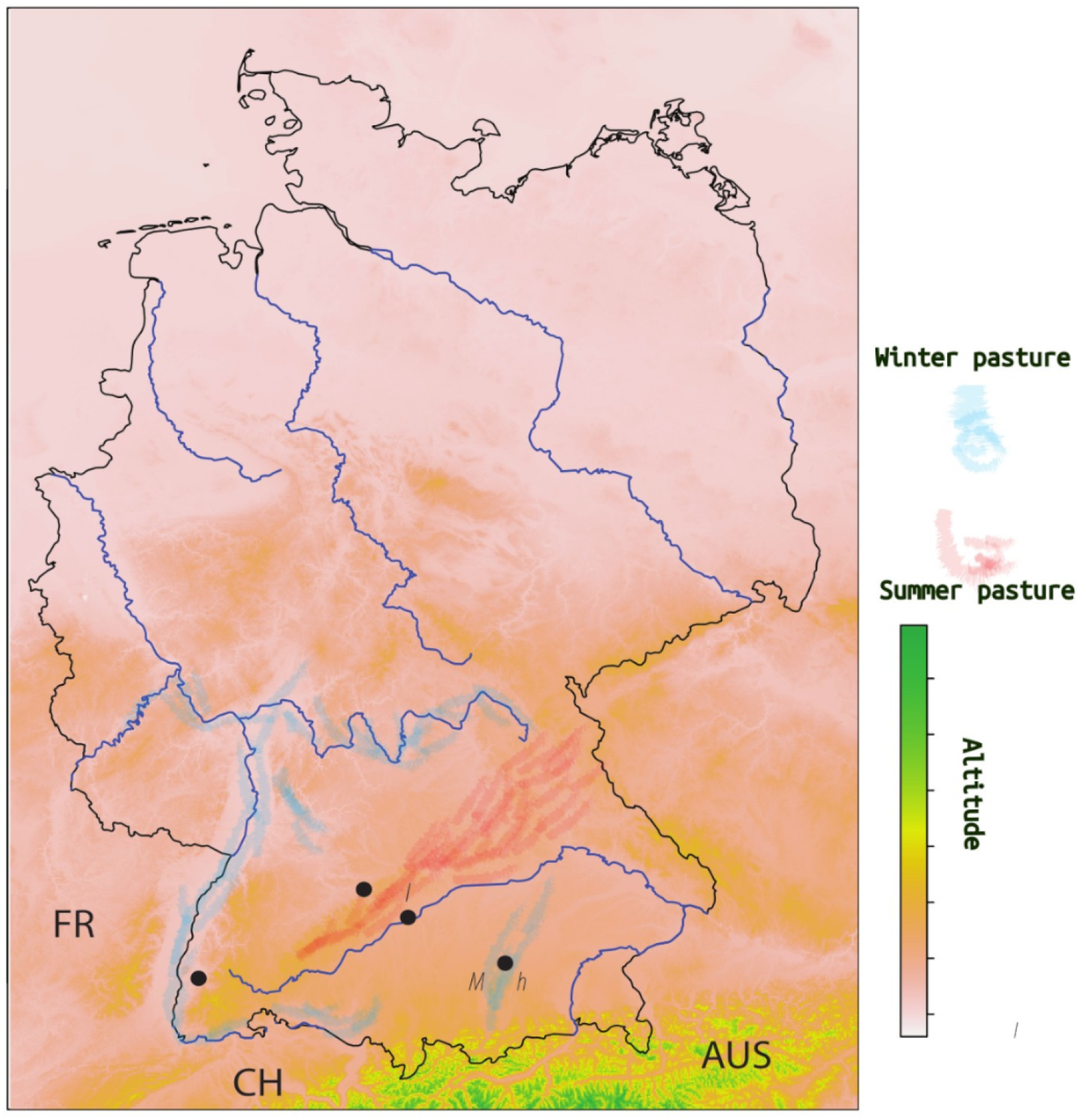
Sketch of the transhumant cycle, illustrating the correspondence between changes in elevation, the pastoral transhumance cycle, and the autumn outbreaks. Caves are suggested to have played a major role as a nexus where wild reservoirs, vectors, and domesticates meet.

In addition to the historical evidence from the well-documented setting of Southern Germany, we can obtain some preliminary support by looking for spatial association between pastoral activities and plague outbreaks. We tested this association in the present using a compilation by World Health Organization of recent (2016) disease foci (62) and data on land use intensity for pastoralism, rangeland, and grazing land for that same year (2016), obtained from the HYDE 3.3 database (63) (Figure 3, A). Random sampling of the pastoral land use intensity inside and outside of plague focus regions (see SI1 for data and scripts) suggests a very strong and significant association between intensive herding and plague foci (Figure 3, B-C).

**Figure 3:**
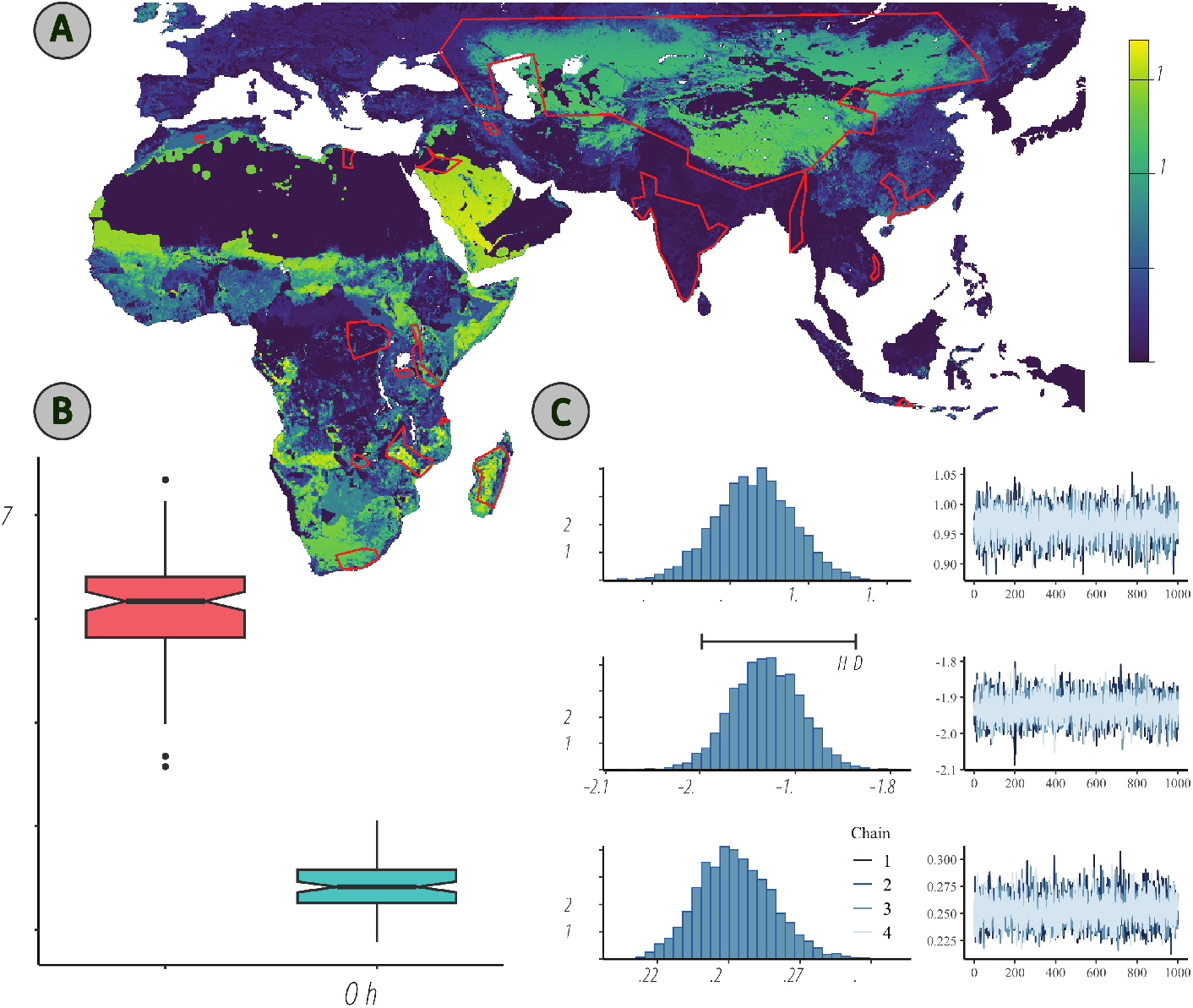
(A) WHO polygons for modern plague distribution and FAO pastoral land use map (colors reflect km2 / cell). (B) comparison of the pixel values inside and outside the polygons, using the map units. The differences are statistically-significant, with strong effect size (One-Way ANOVA p < 0.00001, Eta2=0.92). (C) Bayesian regression parameters based on flat priors, yielding -1.99 to -1.83 95 percent credible intervals for the slope, using standardized data. See SI 1 for further analyses, diagnostics, and script.

We also tested the same association for our study region in Central Europe, using Eckert’s analysis of 16th-17th century parish death records to mark regions where plague outbreaks started in inland regions (64), and utilizing HYDE 3.3 land use data projected to 1600 AD based on population size estimates and land suitability (63) (Figure 4, A). The results show, again, significant association between land use for pasture and inland origins for plague outbreaks (Figure 4, C-D) – although the effect size is not as strong as in the modern dataset.

**Figure 4:**
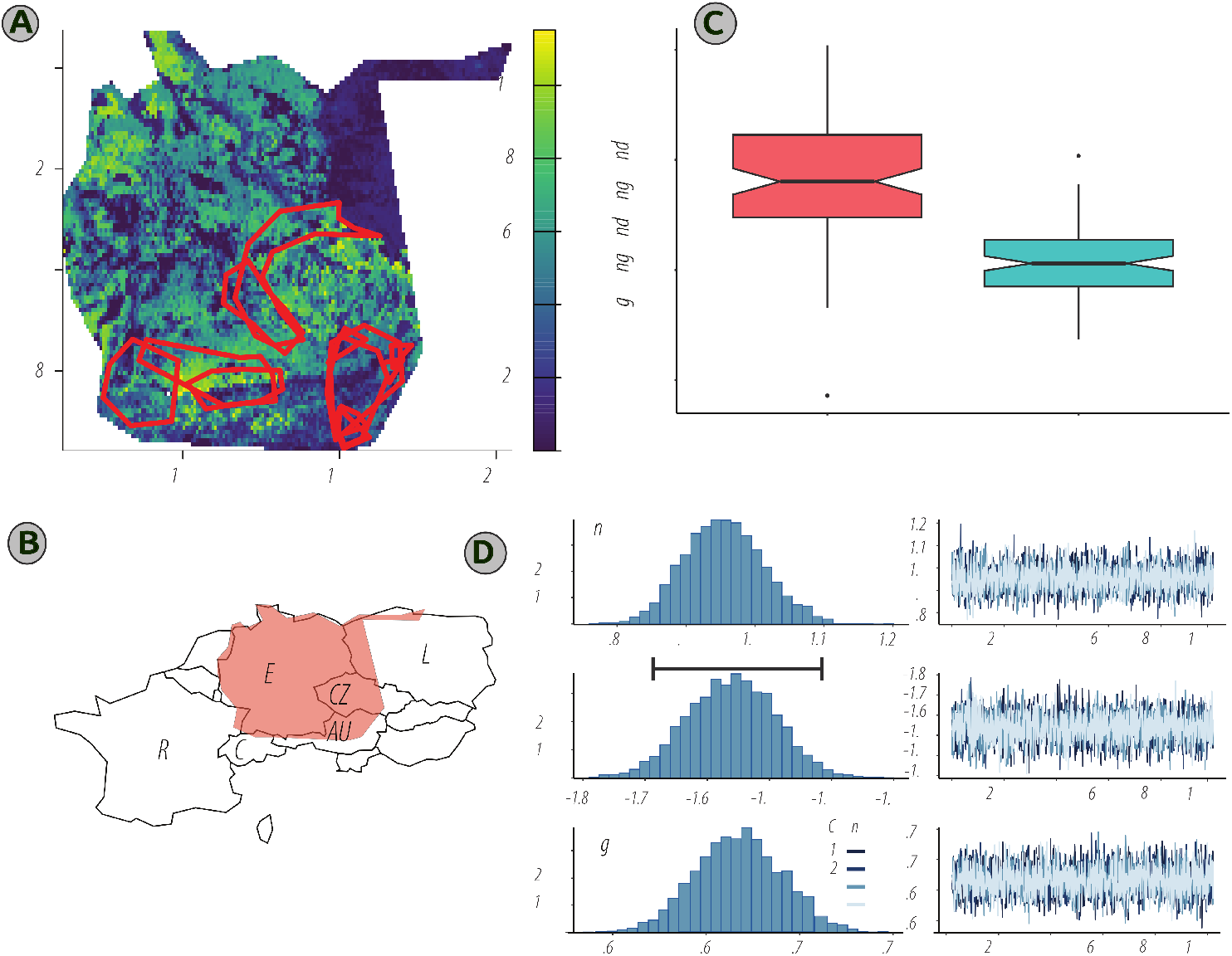
(A) polygons marking first-year records of inland outbreaks in late 16th and 17th centuries Europe, based on Eckert’s (2000) burial record data. The background is HYDE 3.3 pastoral land use map, modelled for 1600 AD (colors reflect km2 / cell). (B) The expanse marked by the polygon encompassing the extreme points of Eckert’s records, which served as a background to the analysis, displayed against a schematic geopolitical map for orientation. (C) comparison of the pixel values inside and outside the polygons, using the map units. The differences are statistically-significant, with strong effect size (One-Way ANOVA p < 0.00001, Eta2=0.38). (D) Bayesian regression parameters based on flat priors, yielding -1.68 to -1.42 95 percent credible intervals for the slope, using standardized data. A prior for the slope was set based on the values of the slope in the modern dataset. See SI 1 for further analyses, diagnostics, and script.

## 3 Discussion

We hypothesize that vertical transhumance in 14th–19th century Europe was responsible for at least some of the plague outbreaks that started in inland, mountainous regions, harbouring plague bacteria. We propose that the specific point of encounter between Y. pestis and humans was through infected sediments and vectors in cave settings, situated at high altitudes potentially associated with plague reservoirs, acting as a hub for inter-specific encounters in marginal settings with karst ecology, and providing stable conditions for the bacteria to survive in local hosts or sediments, as temporary or short-term reservoirs. Furthermore, we consider the potential diversity of livestock herds, comprising – depending on region – of sheep and goat herds and associated dogs, and potentially other domesticates to be a part of the chain of infection under this scenario, acting as intermediate or ‘secondary’ hosts, alias temporary reservoirs that carry the bacteria and fleas from natural foci to human population centers.

The role of nomads in plague transmission chain has been noted in the literature before, but without an explicit mechanism relating the essential connection between transhumance and inland mode of plague spread. The transition from viewing generalised ‘nomads’ as transhumant pastoralists opens new paths to understanding plague dynamics, allowing us to observe the ecology and epidemiology of the disease. These include better appreciation of the potential role of livestock animals and domesticated dogs as temporary, domesticated reservoirs alongside commensal rats, and also the role of seasonal shelters as foci for disease, especially caves in karst regions.

The transhumance biocultural pathway hypothesis also substitutes contingency in encountering plague vectors with a model based on a ubiquitous cultural-ecological phenomenon, which can be empirically falsified. Although the seasonal and spatially-explicit data from early-modern Central Europe provided a fairly simple way to falsification, it is not more than a suggestion due to the problems inherent in the cumulative errors begotten by merging second-order data sources, even Eckert’s excellent geomedical dataset and HYDE’s state-of-the-art modelled land use maps. A conservative interpretation of our evidence is that it does not contradict the transhumance biocultural pathway hypothesis, which awaits further interdisciplinary investigation, based on additional in-depth historical analysis, archaeological fieldwork and genetic lab work.

## 4 Materials and Methods

We compiled two primary datasets to investigate the relationship between plague foci and pastoral land use.

The first dataset, focusing on recent plague occurrences, utilized a 2016 compilation of disease foci from the World Health Organization (62). We manually digitized these WHO map polygons using Google Earth™ and overlaid them onto raster data representing land use intensity for pastoralism, rangeland, and grazing land for the same year (2016). These rasters, obtained from the HYDE 3.3 database (https://hyde-portal.geo.uu.nl/63) at a spatial resolution of 5 arcmin (9.21 x 9.21km), differentiate between pastoral land use (managed land for pasture) and rangeland (unmodified areas used for pastoral activity). For this analysis, however, these categories were merged. The HYDE database, covering 1960 to the present, is based on Food and Agriculture Organization (FAO) data (https://hyde-portal.geo.uu.nl/63).

The second dataset examined historical plague outbreaks. It drew upon Eckert’s analysis of 16th-17th century parish death records in Central Europe, identifying regions where inland plague outbreaks originated (64). We digitized twelve polygons (SI1) from Eckert’s maps to define these historical inland outbreak regions. We then compared the intensity of pastoral land use both inside and outside these polygons, using HYDE 3.3 land use data projected to 1600 AD. This projection was based on population size estimates and land suitability. The background sample, outside the polygons, encompassed the geographical coverage of Eckert’s data, which includes modern Germany and German-speaking parts of present-day Austria, Switzerland, France, Poland, and the Czech Republic. This broad background was chosen to avoid the limitations of modern political boundaries.

For our analysis, we sampled the combined pastoral land-use data rasters (modern and 1600 CE) 100 times within the sub-areas defined by the WHO plague foci polygons. Each iteration provided the mean pixel value from 100 individual sub-samples. Areas of the raster outside the polygons were sampled similarly, with the number of repetitions adjusted for area discrepancies. We then regressed standardized sample values on a dummy variable to assess the effect of group (within or outside plague foci polygons) on the slope. A parallel analysis was conducted using reconstructed land use rasters from 1600 CE (also from the HYDE dataset) and plague foci polygons derived from Eckert (64). These analyses were executed in R 4.4.1, utilizing the ‘terra’ (65), ‘brms’ (66), ‘tidyverse’ (67), ‘geodata’ (68), ‘rnaturalearth’ (69), and ‘effectsize’ (70) packages. Polygons were digitally rendered by NM using Google Earth™. All data files and scripts are available in SI1 and via https://doi.org/10.5281/zenodo.15680661.

## 5 Acknowledgments

The work was supported by the ERC Synergy (101118880)/UKRI EPSRC Horizon Europe UK Guarantee Funding (EP/Z003288/1) ‘Synergy-Plague’ Grant.

## 6 Supplements

Dataset S1 (separate file – available at https://doi.org/10.5281/zenodo.15680661) including the scripts, output files.

